# Mathematical constraints on *F*_*ST*_: multiallelic markers in arbitrarily many populations

**DOI:** 10.1101/2021.07.23.453474

**Authors:** Nicolas Alcala, Noah A. Rosenberg

## Abstract

Interpretations of values of the *F*_*ST*_ measure of genetic differentiation rely on an understanding of its mathematical constraints. Previously, it has been shown that *F*_*ST*_ values computed from a biallelic locus in a set of multiple populations and *F*_*ST*_ values computed from a multiallelic locus in a pair of populations are mathematically constrained as a function of the frequency of the allele that is most frequent across populations. We generalize from these cases to report here the mathematical constraint on *F*_*ST*_ given the frequency *M* of the most frequent allele at a *multiallelic* locus in a set of *multiple* populations. Using coalescent simulations of an island model of migration with an infinitely-many-alleles mutation model, we argue that the joint distribution of *F*_*ST*_ and *M* helps in disentangling the separate influences of mutation and migration on *F*_*ST*_. Finally, we show that our results explain a puzzling pattern of microsatellite differentiation: the lower *F*_*ST*_ in an interspecific comparison between humans and chimpanzees than in the comparison of chimpanzee populations. We discuss the implications of our results for the use of *F*_*ST*_.

## 1. Introduction

Multiallelic loci such as microsatellites and haplotype assignments are used to study genetic differentiation in a variety of fields, ranging from ecology and conservation genetics to anthropology and human genomics. Genetic differentiation is often measured for multiallelic loci using the multiallelic extension of Wright’s fixation index *F*_*ST*_ [1]:

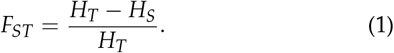

For a polymorphic multiallelic locus with *I* distinct alleles in a set of *K* subpopulations, denoting by *p*_*k,i*_ the frequency of allele *i* in subpopulation 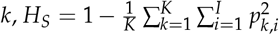 and 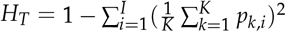.

*F*_*ST*_ values are known to be smaller for multiallelic than for biallelic loci [2]. One reason invoked to explain this difference is that within-subpopulation heterozygosity *H*_*S*_ mathematically constrains the maximal value of *F*_*ST*_ to be below 1, and the constraint is stronger when *H*_*S*_ is high. This phenomenon was noticed concurrently in simulation-based, empirical, and theoretical studies [3, 4, 5, 6, 7], and the mathematical constraints describing the dependence were subsequently clarified [8, 9].

Studies have found that the maximal value of *F*_*ST*_ can be viewed as constrained not only by functions of the within-subpopulation allele frequency distribution such as *H*_*S*_, but alternatively by aspects of the global allele frequency distribution across subpopulations. For a biallelic locus in *K* = 2 subpopulations, Maruki *et al*. [10] showed that the maximal *F*_*ST*_ as a function of the frequency *M* of the most frequent allele decreases as *M* increases from 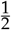 to 1 (see also [11]). Generalizing the biallelic case to arbitrarily many alleles, Jakobsson *et al*. [12] showed that for multiallelic loci with an unspecified number of distinct alleles, the maximal *F*_*ST*_ increases from 0 to 1 as a function of *M* if 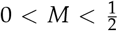, and decreases from 1 to 0 for 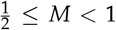 in the manner reported by Maruki *et al*. [10] for biallelic loci. Edge and Rosenberg [13] generalized these results to the case of a fixed finite number of alleles, showing that the maximal *F*_*ST*_ differs slightly from the unspecified case when the fixed number of distinct alleles is an odd number.

Generalizing the simplest case of *K* = *I* = 2 in a different direction, Alcala and Rosenberg [14] considered biallelic loci in the case of a fixed number of subpopulations *K* ≥ 2. We showed that the maximal value of *F*_*ST*_ displays a peculiar behavior as a function of *M*: the upper bound has a maximum of 1 if and only if 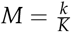, for integers *k* with 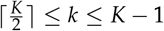. The constraints on the maximal value of *F*_*ST*_ dissipate as *K* tends to infinity, even though for any fixed *K*, there always exists a value of *M* for which 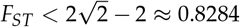.

Relating *F*_*ST*_ to its maximum as a function of *M* helps explain surprising phenomena that arise during population-genetic data analysis. For example, Jakobsson *et al*. [12] showed that stronger constraints on *F*_*ST*_ could explain the low *F*_*ST*_ values seen in pairs of African human populations. They also found that such constraints could explain the lower *F*_*ST*_ values seen in high-diversity multiallelic loci compared to lower-diversity loci—microsatellites compared to single-nucleotide polymorphisms. Alcala and Rosenberg [14] showed that constraints on the maximal *F*_*ST*_ could explain the lower *F*_*ST*_ values between human populations seen when computing *F*_*ST*_ pairwise rather than from all populations simultaneously.

In this study, we characterize the relationship between *F*_*ST*_ and the frequency *M* of the most frequent allele, for a *multiallelic* locus and an arbitrary specified value of the number of subpopulations *K*. We derive the mathematical upper bound on *F*_*ST*_ in terms of *M*, extending the biallelic result of Alcala and Rosenberg [14] to the multiallelic case, and providing the most comprehensive description of the mathematical constraints on *F*_*ST*_ in terms of *M* to date (Table 1). To assist in interpreting the new bound, we simulate the joint distribution of *F*_*ST*_ and *M* in the island migration model, describing its properties as a function of the number of subpopulations, the migration rate, and a mutation rate. The *K*-subpopulation upper bound on *F*_*ST*_ in terms of *M* facilitates an explanation of counterintuitive aspects of inter-species genetic differentiation. We discuss the importance of the results for applications of *F*_ST_ more generally.

**Table 1.** Studies describing the mathematical constraints on *F*_*ST*_.

| Reference | Number of alleles | Number of subpopulations | Variable in terms of which constraints are reported* |
| --- | --- | --- | --- |
| LONG and KITTLES [8] | unspecified value $\geq 2$ | fixed finite value $\geq 2$ | $H_S$ |
| ROSENBERG *et al.* [11] | 2 | 2 | $\delta$ |
| HEDRICK [9] | unspecified value $\geq 2$ | fixed finite value $\geq 2$ | $H_S$ |
| MARUKI *et al.* [10] | 2 | 2 | $H_S, M$ |
| JAKOBSSON *et al.* [12] | unspecified value $\geq 2$ | 2 | $H_T, M$ |
| EDGE and ROSENBERG [13] | fixed finite value $\geq 2$ | 2 | $H_T, M$ |
| ALCALA and ROSENBERG [14] | 2 | fixed finite value $\geq 2$ | $M$ |
| This paper | unspecified value $\geq 2$ | fixed finite value $\geq 2$ | $M$ |

$H_S$ and $H_T$ denote the within-subpopulation and total heterozygosities, respectively. $\delta$ denotes the absolute difference in the frequency of a specific allele between two subpopulations, and $M$ denotes the frequency of the most frequent allele in the total population. Instead of heterozygosities $H_S$ or $H_T$ , some studies consider homozygosities $1 - H_S$ or $1 - H_T$ .

## 2. Model

Our goal is to derive the range of values *F*_*ST*_ can take—the lower and upper bounds on *F*_*ST*_ —as a function of the frequency *M* of the most frequent allele for a multiallelic locus, when the number of subpopulations *K* is a fixed finite value greater than or equal to 2. We follow previous studies [12, 13, 14, 15] in describing notation and constructing the scenario.

We consider a polymorphic locus with an unspecified number of distinct alleles, in a setting with *K* subpopulations contributing equally to the total population. We denote the frequency of allele *i* in subpopulation *k* by *p*_*k,i*_, with sum 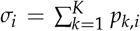 across subpopulations. Each allele frequency *p*_*k,i*_ lies in [0, 1]. Within subpopulations, allele frequencies sum to 1: for each 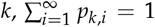. Hence, *σ*_*i*_ lies in [0, *K*], and 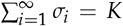. We number alleles from most to least frequent, so *σ*_*i*_ ≥ *σ*_*j*_ for *i* ≤ *j*.

Because by assumption the locus is polymorphic, *σ*_*i*_ < *K* for each *i*. Alleles 1 and 2 have nonzero frequency in at least one subpopulation, not necessarily the same one; we have *σ*_1_ > 0 and *σ*_2_ > 0. We denote the mean frequency of the most frequent allele across subpopulations by *M* = *σ*_1_/*K*. We then have 0 < *M* < 1. We treat the allele frequencies *p*_*k,i*_ and associated quantities *M* and *σ*_*i*_ as parametric values, and not as estimates computed from data.

Eq. 1 expresses *F*_*ST*_ as a ratio involving within-subpopulation heterozygosity, *H*_*S*_, and total heterozygosity, *H*_*T*_, with 0 ≤ *H*_*S*_ < 1 and 0 ≤ *H*_*T*_ < 1. Because we assume the locus is polymorphic, *H*_*T*_ > 0. We write eq. 1 in terms of allele frequencies, permitting the number of distinct alleles to be arbitrarily large:

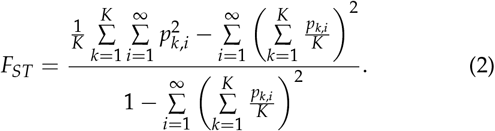

Hence, our goal is, for fixed *σ*_1_ = *KM*, 0 < *σ*_1_ < *K*, to identify the matrices (*p*_*k,i*_)_*K×*∞_, with *p*_*k,i*_ in [0, 1], 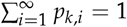 and 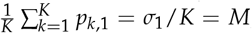, that minimize and maximize *F*_*ST*_ in eq. 2.

Note that we adopt the interpretation of *F*_*ST*_ as a “statistic” that describes a mathematical function of allele frequencies rather than as a “parameter” that describes coancestry of individuals in a population [e.g. 16]. See Alcala and Rosenberg [14] for a discussion of interpretations of *F*_*ST*_ when studying its mathematical properties.

## 3. Mathematical constraints

### (a) Lower bound of *F*_*ST*_

Bounds on *F*_*ST*_ in terms of the frequency of the most frequent allele can be written with respect to *M* or *σ*_1_, noting that *M* ranges in (0, 1) and *σ*_1_ ranges in (0, *K*). For the lower bound, from eq. 2, for any choice of *σ*_1_, *F*_*ST*_ = 0 can be achieved. Consider (*σ*_1_, *σ*_2_, …) with *σ*_*i*_ in [0, *K*) for each *k, σ*_*i*_ ≥ *σ*_*j*_ for *i* ≤ *j*, 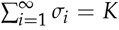, and *σ*_1_ > 0 and *σ*_2_ > 0. We set *p*_*k,i*_ = *σ*_*i*_ /*K* for all subpopulations *k* and alleles *i*; this choice yields *F*_*ST*_ = 0.

*F*_*ST*_ = 0 implies that the numerator of eq. 2, *H*_*T*_ − *H*_*S*_, is zero. This numerator can be written 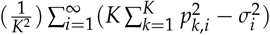. The Cauchy-Schwarz inequality guarantees that 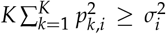, with equality if and only if *p*_1_,_*i*_ *= p*_*2,i*_ =… = *p*_*k,i*_ = *σ*_*i*_/*K*. Applying the Cauchy-Schwarz inequality to all alleles *i*, the numerator of eq. 2 is zero only if for all *i*, (*p*_1,*i*_, *p*_2,*i*_, …, *p*_*K,i*_) = (*σ*_*i*_ /*K, σ*_*i*_ /*K*, …, *σ*_*i*_ /*K*).

Thus, we can conclude that the allele frequency matrices in which all *K* subpopulations have identical allele frequency vectors are the only matrices for which *F*_*ST*_ = 0. The lower bound on *F*_*ST*_ is equal to 0 irrespective of *M* or *σ*_1_, for any value of the number of subpopulations *K*.

### (b) Upper bound of *F*_*ST*_

To derive the upper bound on *F*_*ST*_ in terms of *M* = *σ*_1_/*K*, we must maximize *F*_*ST*_ in eq. 2, assuming that *σ*_1_ and *K* are constant. The computations are performed in the Appendix; we write the main result as a function of *σ*_1_, noting that it can be converted into a function of *M* by replacing *σ*_1_ with *KM*.

In Theorem 1, we treat the case in which *σ*_1_ has an integer value. For non-integer *σ*_1_, Theorem 2 shows that the maximal *F*_*ST*_ requires that (i) the sum of squared allele frequencies across alleles and subpopulations, 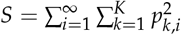, is maximal, and (ii) alleles *i* = 2, 3, … are each present in at most one subpopulation, but allele 1 might be present in more than one subpopulation. We then separately maximize *F*_*ST*_ as a function of *σ*_1_ for *σ*_1_ in (0, 1) and non-integer *σ*_1_ in (1, *K*). These two cases differ in that allele 1 appears in a single subpopulation in the former case, and it must appear in at least two subpopulations in the latter.

The maximal *F*_*ST*_ as a function of *σ*_1_ for *σ*_1_ in (0, *K*) is

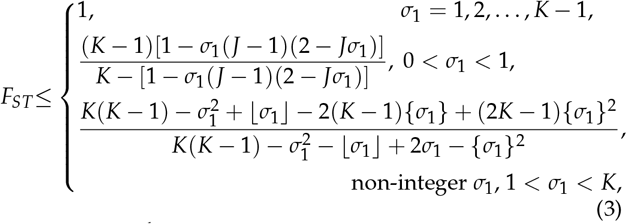

where 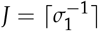. Here, ⌈*x*⌉ denotes the smallest integer greater than or equal to *x*, ⌊*x*⌋ denotes the greatest integer less than or equal to *x*, and {*x*} = *x*−⌊*x*⌋ denotes the fractional part of *x*. Note that for an integer choice of *σ*_1_, the maximum from eq. 3 and the limits as *σ*_1_ tends to the integer from above and below all equal 1, so that the maximum as a function of *σ*_1_ is continuous.

From the Appendix, *F*_*ST*_ reaches its upper bound for integer *σ*_1_ when allele 1 has frequency 1 in each of *σ*_1_ subpopulations, and when in each of the remaining *K*− *σ*_1_ subpopulations, an allele other than allele 1 has frequency 1. These alleles of frequency 1 need not be private, although they can be; any identity relationships among them are permissible, provided that when summing frequencies across subpopulations, none of these alleles has a sum that exceeds *σ*_1_. The locus can have as few as 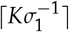 alleles of nonzero frequency and as many as *K* − *σ*_1_ + 1.

For *σ*_1_ in interval (0, 1), *F*_*ST*_ is maximal when each allele is present in only a single subpopulation, and when each subpopulation has exactly *J* alleles with a nonzero frequency: *J* − 1 alleles at frequency *σ*_1_ and one allele at frequency 1 − (*J* − 1)*σ*_1_ ≤ *σ*_1_. Because each subpopulation has *J* distinct alleles and no alleles are shared across subpopulations, this upper bound requires that the locus has *KJ* alleles of nonzero frequency.

For non-integer *σ*_1_ in (1, *K*), *F*_*ST*_ reaches its maximum when there are ⌊*σ*_1_⌋ subpopulations in which the most frequent allele has frequency 1, a single subpopulation in which it has frequency {*σ*_1_} and a private allele has frequency 1 − {*σ*_1_}, and *K*− ⌊*σ*_1_⌋ − 1 subpopulations each with a different private allele at frequency 1. Only the most frequent allele is shared across subpopulations, and a single subpopulation displays polymorphism. At the maximum, *K* − ⌊*σ*_1_⌋ + 1 alleles have nonzero frequency.

### (c) Properties of the upper bound

Figure 1 shows the maximal value of *F*_*ST*_ in terms of *M* = *σ*_1_/*K* for various values of the number of subpopulations, *K*. We describe a number of properties of this upper bound.

**Figure 1.**
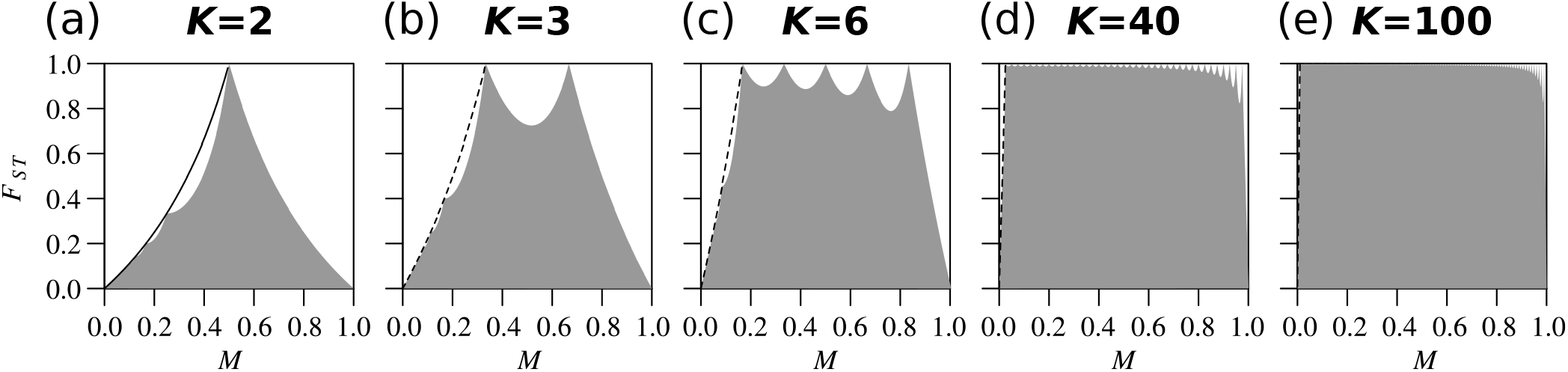
Bounds on *F*_*ST*_ as a function of the frequency of the most frequent allele, *M*, for a multiallelic locus, for each of several different numbers of subpopulations *K*. (a) *K* = 2. (b) *K* = 3. (c) *K* = 6. (d) *K* = 40. (e) *K* = 100. The gray region represents the space between the upper and lower bounds on *F*_*ST*_. The dashed line represents the curve that the jagged maximal *F*_*ST*_ touches when 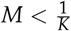, computed from eq. 4. The upper bound is computed from eq. 3; for each *K*, the lower bound is 0 for all values of *M*.

#### Piecewise structure of the upper bound

First, we observe that the upper bound has a piecewise structure.

For 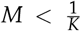, the upper bound depends on 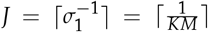. As *KM* increases in (0, 1), each decrement in the integer value of 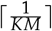 produces a distinct “piece” with domain 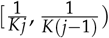, for integers *j* ≥ 2. Within each interval 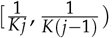, *J* has the constant value *j*.

At 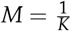, the upper bound has its first transition between cases. For 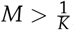, the upper bound depends on ⌊*σ*_1_⌋ = ⌊*KM*⌋. As *KM* increases in [1, *K*), each increment in ⌊*KM*⌋ also produces a distinct piece of the domain. For each *k* from 1 to *K* − 1, ⌊*KM*⌋ = *k* for *M* in 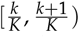.

Counting the intervals of the domain, we see that an infinite number of distinct intervals occur for *M* in 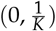, and *K* − 1 intervals occur for *M* in 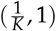. Within intervals, the function describing the upper bound is smooth.

#### Behavior of the upper bound for 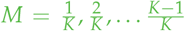

The upper bound is equal to 1 at 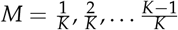. For *M* in 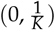, setting the numerator and denominator equal in eq. 3, we find that the upper bound is never equal to 1. For *M* in 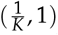, the upper bound is equal to 1 if and only if {*σ*_1_} = 0, that is, if and only if *σ*_1_ is an integer and 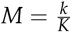 for *k* = 2, 3, …, *K* − 1.

Hence, noting that the upper bound is equal to 1 at 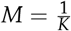, we conclude that the upper bound can equal 1 if and only if 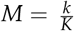 for integers *k* = 1, 2, …, *K* − 1. For fixed *K*, the upper bound on *F*_*ST*_ has exactly *K* − 1 maxima at which *F*_*ST*_ can equal 1, at 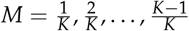. We can conclude that *F*_*ST*_ is unconstrained within the unit interval only for a finite set of values of the frequency *M* of the most frequent allele. The size of this set increases with the number of subpopulations *K*.

#### Behavior of the upper bound for *M* in 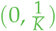

For *M* in 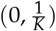, we can compute the value of the upper bound at the transition points between distinct pieces of the domain, namely values of 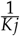 for integers *j* ≥ 2. Applying eq. 3, we observe that at 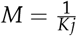, the upper bound has value 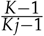. In other words, the upper bound touches the curve

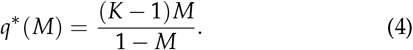

This curve is represented in Fig. 1 as a dashed line.

Note that for *K* = 2, the special case considered by Jakobsson *et al*. [12], eq. 4 reduces to *q*\* (*M*) = *M*/(1 − *M*) = *σ*_1_/(2 − *σ*_1_), which matches eq. 21 from Jakobsson *et al*. [12]. In fact, setting *K* = 2, eq. 3 for *M* in 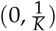 reduces to the *K* = 2 upper bound on *F*_*ST*_ in eq. 9 of [12].

#### Behavior of the upper bound for *M* in 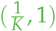

Because the upper bound is a smooth function on each interval of its domain, and because it possesses maxima at interval boundaries 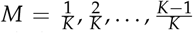, it must possess local minima in intervals 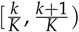 for *k* = 1, 2, …, *K* − 2. Indeed, such minima are visible in Figure 1 in cases with *K* = 3, *K* = 6, *K* = 40, and *K* = 100; for *K* = 2, only one maximum occurs, so that there is no interval between a pair of maxima in which a minimum can occur. Note that because we restrict attention to *M* in (0, 1), we do not count the point at *M* = 1 and *F*_*ST*_ = 0 as a local minimum.

## 4. Joint distribution of *M* and *F*_*ST*_ under an evolutionary model

So far, we have described the mathematical constraint imposed on *F*_*ST*_ by *M* without respect to the frequency with which particular values of *M* arise in evolutionary scenarios. As an assessment of the bounds in evolutionary models can illuminate the settings in which they are most salient in population-genetic data analysis [9, 14, 17, 18, 19, 20], we simulated the joint distribution of *F*_*ST*_ and *M* under an island migration model, relating the distribution to the mathematical bounds on *F*_*ST*_. This analysis considers allele frequency distributions, and hence values of *M* and *F*_*ST*_, generated by evolutionary models. The simulation approach is modified from [14, 15].

### (a) Simulations

We simulated alleles under a coalescent model, using the software `MS` [21]. We considered a total population of *KN* diploid individuals subdivided into *K* subpopulations of size *N*. At each generation, a proportion *m* of the individuals in a subpopulation originated outside the subpopulation. Thus, the scaled migration rate is 4*Nm*, and it corresponds to twice the number of individuals in a subpopulation that originate elsewhere. We considered the island model [22, 23, 24], in which migrants have the same probability 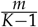 to come from any specific other subpopulation. We used an infinitely-many-alleles model; mutations occur at rate *μ*, and the scaled mutation rate is 4*Nμ*.

We examined three values of *K* (2, 6, 40), three values of 4*Nμ* (0.1, 1, 10), and three values of 4*Nm* (0.1, 1, 10). Note that in `MS`, time is scaled in units of 4*N* generations, and there is no need to specify subpopulation sizes *N*. `MS` simulates an infinitely-many-sites model, where each mutation occurs at a new site; each haplotype is a new allele, so that each mutation creates a new allele. For our analysis, we are concerned only with the allelic categories, and not with the simulated sequences; thus, although the simulation follows the infinitely-many-sites model, the analysis treats simulated data sets as having been generated under an infinitely-many-alleles model.

For each parameter triplet (*K*, 4*Nμ*, 4*Nm*), we performed 1,000 replicate simulations, sampling 100 sequences per subpopulation in each replicate. We computed *F*_*ST*_ values from the parametric allele (haplotype) frequencies. `MS` commands appear in File S1; note that the simulation approach here uses the standard method of simulating `MS` with a specified mutation rate *θ* = 4*Nμ*, whereas in our previous analyses of biallelic cases [14, 15], we had employed the alternative approach of requiring simulated datasets to possess exactly one segregating site.

Figure 2 shows the joint distribution of *M* and *F*_*ST*_ for the nine values of (4*Nμ*, 4*Nm*) in the case of *K* = 2. Figures S1 and S2 provide similar figures for *K* = 6 and *K* = 40, respectively.

**Figure 2.**
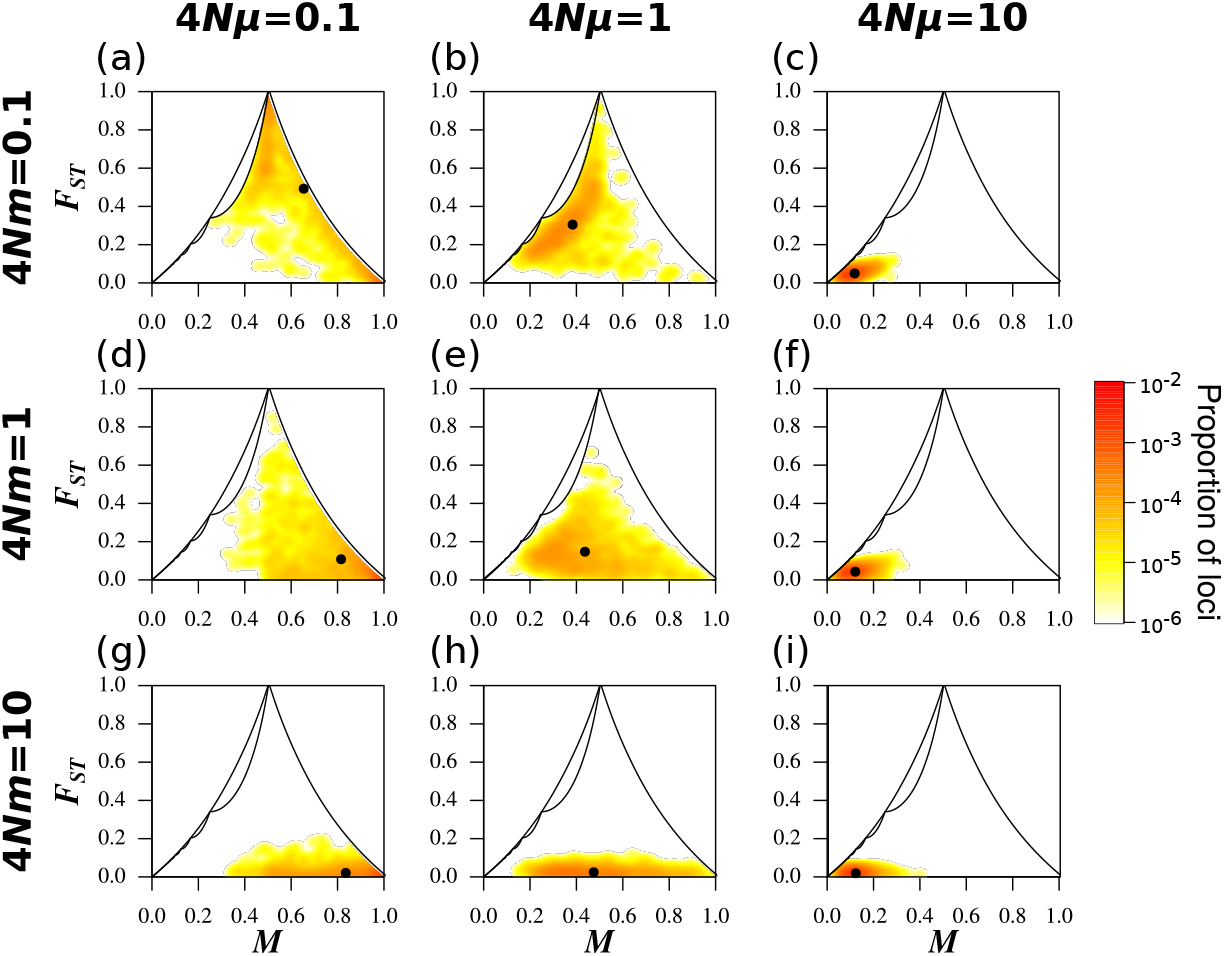
Joint density of the frequency *M* of the most frequent allele and *F*_*ST*_ in the island migration model with *K* = 2 subpopulations, for different scaled migration rates 4*Nm* and mutation rates 4*Nμ*. (a) 4*Nμ* = 0.1, 4*Nm* = 0.1. (b) 4*Nμ* = 1, 4*Nm* = 0.1. (c) 4*Nμ* = 10, 4*Nm* = 0.1. (d) 4*Nμ* = 0.1, 4*Nm* = 1. (e) 4*Nμ* = 1, 4*Nm* = 1. (f) 4*Nμ* = 10, 4*Nm* = 1. (g) 4*Nμ* = 0.1, 4*Nm* = 10. (h) 4*Nμ* = 1, 4*Nm* = 10. (i) 4*Nμ* = 10, 4*Nm* = 10. The black solid line represents the upper bound on *F*_*ST*_ in terms of *M* (eq. 3); the black point plots the mean values of *M* and *F*_*ST*_. Colors represent the density of loci, estimated using a Gaussian kernel density estimate with a bandwidth of 0.02, with density set to 0 outside of the bounds. Loci are simulated using coalescent software `MS`, assuming an island model of migration and an infinitely-many-alleles mutation model. Each panel considers 1,000 replicate simulations, with 100 lineages sampled per subpopulation. Figures S1 and S2 present similar results for *K* = 6 and *K* = 40 subpopulations, respectively.

### (b) Impact of the mutation rate

For fixed migration rate 4*Nm* and number of subpopulations *K*, the main impact of the mutation rate is on the frequency *M* of the most frequent allele. For *K* = 2, under weak mutation (4*Nμ* = 0.1), the joint distribution of *M* and *F*_*ST*_ is highest in the high-*M* region, for all values of 4*Nm* (Fig. 2A, D, G). Although most simulation replicates produce 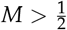 with an upper bound on *F*_*ST*_ less than one, this set of parameter values does give rise to replicates near the peak at 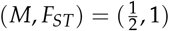.

Under intermediate mutation (4*Nμ* = 1), the increased mutation rate tends to decrease *M*, shifting the joint distribution to lower values of *M* for all values of 4*Nm* (Fig. 2B, E, H). Finally, under strong mutation (4*Nμ* = 10), the joint distribution of *M* and *F*_*ST*_ is highest in the low-*M* region, for all values of 4*Nm* (Fig. 2C, F, I). In this region, the upper bound on *F*_*ST*_ is most strongly constrained, leading to low *F*_*ST*_ values.

### (c) Impact of the migration rate

For fixed mutation rate 4*Nμ* and number of subpopulations *K*, the impact of the migration rate is seen primarily in the *F*_*ST*_ values rather than the values of *M*. Under weak migration (4*Nm* = 0.1), subpopulations are differentiated, and the joint distribution of *M* and *F*_*ST*_ is highest near the upper bound on *F*_*ST*_ in terms of *M* (Fig. 2A, B, C).

Under intermediate migration (4*Nm* = 1), differentiation between subpopulations decreases, and the joint density of *M* and *F*_*ST*_ is highest at lower values of *F*_*ST*_ (Fig. 2D, E, F). Under strong migration (4*Nm* = 10), the joint density of *M* and *F*_*ST*_ nears the lower bound (Fig. 2G, H, I).

### (d) Impact of the number of subpopulations

In Figure 1, the number of subpopulations changes the shape of the region in which *F*_*ST*_ is permitted to range as a function of *M*. Thus, in simulations, the impact of the number of subpopulations *K* is observed in cases in which a change in *K* permits *F*_*ST*_ to expand its range within the unit square for (*M, F*_*ST*_). For each of the nine choices of (4*Nμ*, 4*Nm*), Figure 3 summarizes the means observed for (*M, F*_*ST*_) in Figures 2, S1, and S2, corresponding to *K* = 2, *K* = 6, and *K* = 40, respectively.

**Figure 3.**
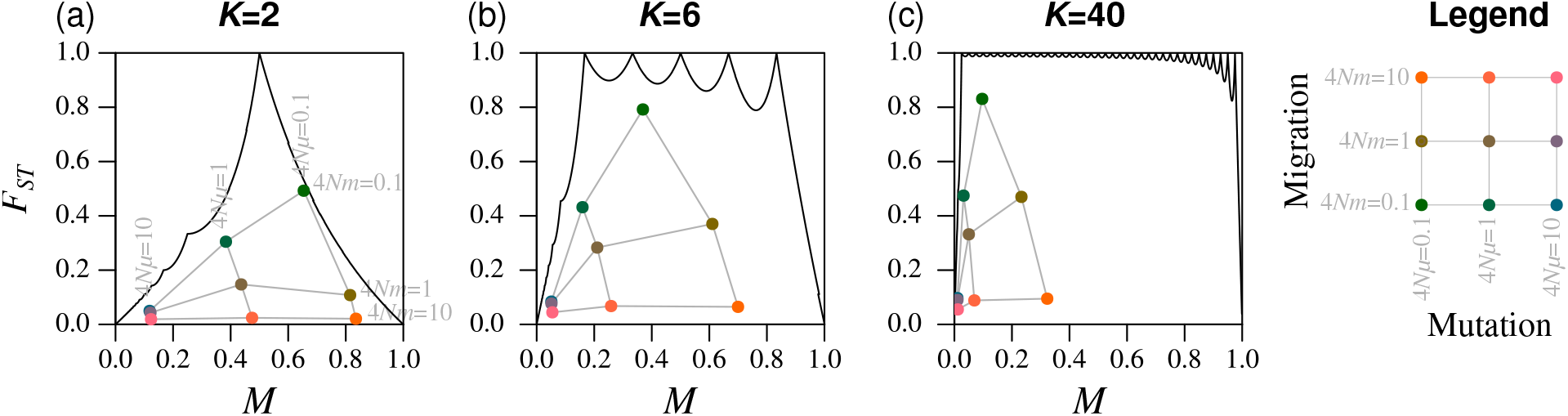
Mean frequency *M* of the most frequent allele and mean *F*_*ST*_ in the island migration model, for different scaled migration rates 4*Nm* and mutation rates 4*Nμ* and different numbers of subpopulations *K*. (a) *K* = 2. (b) *K* = 6. (c) *K* = 40. The black solid line represents the upper bound on *F*_*ST*_ in terms of *M* (eq. 3). The colored points represent the mean *M* and mean *F*_*ST*_, where colors correspond to values of 4*Nm*. These points are taken from Figures 2, S1, and S2.

The number of subpopulations generally increases *F*_*ST*_ for fixed 4*Nμ* and 4*Nm*. For example, the mean *F*_*ST*_ can be substantially larger for *K* = 6 than for *K* = 2. Consider (4*Nμ*, 4*Nm*) = (0.1, 0.1). For *K* = 2, the mean *F*_*ST*_ is near its upper bound (Fig. 3A); for *K* = 6, *F*_*ST*_ is not as close to the bound (Fig. 3B). However, because the upper bound for *K* = 6 exceeds that for *K* = 2, the mean *F*_*ST*_ is nevertheless larger in the case of *K* = 6.

## 5. Example: humans and chimpanzees

We now use our theoretical results to examine genetic differentiation in humans and chimpanzees. Because humans and chimpanzees are distinct species, we might expect a genetic differentiation measure such as *F*_*ST*_ to produce a greater value for a computation between them than for a computation among populations within one or the other. Indeed, studies of multiallelic loci do find that adding chimpanzees to data on multiple human populations increases the value of *F*_*ST*_ [8, 25]. However, we will see that *F*_*ST*_ has a more subtle pattern when considering data on multiple *chimpanzee* populations, and that our theoretical computations explain a surprising result.

We examine data on 246 multiallelic microsatellite loci assembled by Pemberton *et al*. [26] from several studies of worldwide human populations and a study of chimpanzees [27]. We consider *F*_*ST*_ comparisons both between humans and chimpanzees and among populations of chimpanzees. For the human data, we consider all 5795 individuals in the dataset, and for the chimpanzee data, we consider 84 chimpanzee individuals from 6 populations: one bonobo population, and 5 common chimpanzee populations (Central, Eastern, Western, hybrid, and captive).

In the data analysis, we perform a computation to summarize the relationship of *F*_*ST*_ to the upper bound. For a set of *Z* loci, denote by *F*_*z*_ and *M*_*z*_ the values of *F*_*ST*_ and *M* at locus *z*. The mean *F*_*ST*_ for the set, or 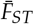, is

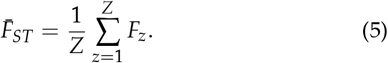

Using eq. 3, we can compute the corresponding maximum *F*_*ST*_ given the observed *σ*_*z*_ = *KM*_*z*_, *z* = 1, 2, …, *Z*. Denoting this quantity by *F*_max,*z*_, we have

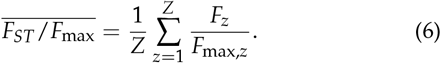

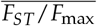 measures the proximity of the *F*_*ST*_ values to their upper bounds: it ranges from 0, if *F*_*ST*_ values at all loci equal 0, to 1, if *F*_*ST*_ values at all loci equal their upper bounds.

We computed the parametric allele frequencies for each subpopulation—the human and chimpanzee groups for the human–chimpanzee comparison, and chimpanzee subpopulations for the comparison of chimpanzees—averaging across subpopulations to obtain the frequency *M* of the most frequent allele. We then computed *F*_*ST*_ and the associated upper bound for each locus, averaging across loci to obtain the overall 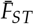 and 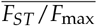 for the full microsatellite set (eqs. 5 and 6).

Surprisingly, given the longer evolutionary time between humans and chimpanzees than among chimpanzee populations, the *F*_*ST*_ value is significantly greater when comparing chimpanzee populations 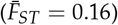 than when comparing humans and chimpanzees (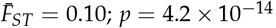, Wilcoxon rank sum test). The explanation for this result can be found in the properties of the upper bound on *F*_*ST*_ given *M*.

Values of *M* are similar in the two comparisons (Fig. 4A, 4B). However, *K* differs, equaling 2 for the human–chimpanzee comparison and 6 for the comparison of chimpanzee subpopulations. Because the theoretical range of *F*_*ST*_ is seen to be smaller for *F*_*ST*_ values computed among smaller sets of subpopulations than among larger sets (Fig. 1), the *F*_*ST*_ values among chimpanzees possess a larger range. For example, the maximal *F*_*ST*_ at the mean *M* of 0.27 observed in pairwise comparisons is 0.34 for *K* = 2 (red segment in Figure 4A), whereas the maximal *F*_*ST*_ at the mean *M* of 0.36 observed for six chimpanzee populations is 0.93 for *K* = 6 (Fig. 4B). Given the stronger constraint in pairwise calculations than in calculations with more subpopulations, it is not unexpected that pairwise *F*_*ST*_ values would be smaller than those in a 6-region computation. A high *F*_*ST*_ among chimpanzees compared to between humans and chimpanzees is a byproduct of mathematical constraints on *F*_*ST*_.

**Figure 4.**
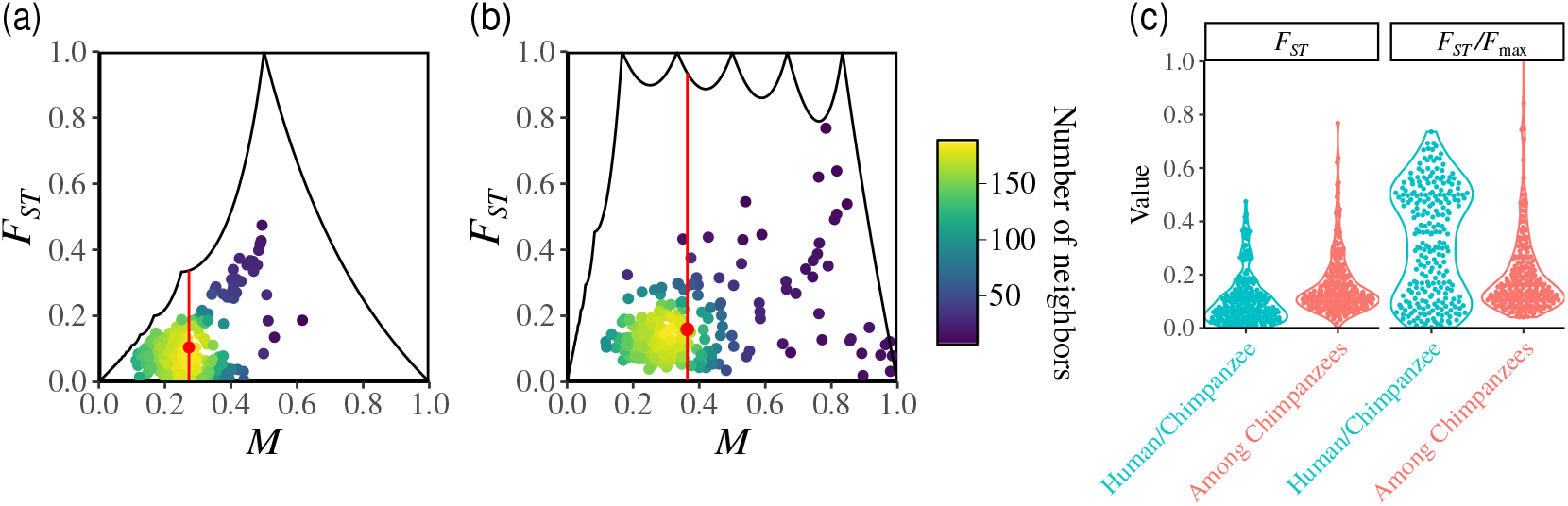
*F*_*ST*_ values for comparisons involving humans and chimpanzees based on multiallelic microsatellite loci. (a) *F*_*ST*_ between humans and chimpanzees, considering *K* = 2 subpopulations (humans, chimpanzees). (b) *F*_*ST*_ among *K* = 6 chimpanzee subpopulations. In (a) and (b), colors represent the number of points in a neighborhood of radius 0.03; red points indicate the mean *M* and *F*_*ST*_, and vertical red segments indicate the permissible range of *F*_*ST*_ at the mean *M*. (c) *F*_*ST*_, computed using eq. 2, and *F*_*ST*_ /*F*_max_, computed using eqs. 2 and 3. Each point plotted represents one locus.

Interestingly, the effect of *K* on *F*_*ST*_ is largely eliminated when each *F*_*ST*_ value is normalized by the associated maximum given *K* and *M* (Fig. 4C). The normalization leads to higher values for human–chimpanzee comparisons than among chimpanzee subpopulations 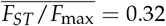 and 0.20, respectively; *p* = 1.1 *×* 10^−9^, Wilcoxon rank sum test), as expected from the greater evolutionary distance between humans and chimpanzees compared to that among chimpanzees.

## 6. Discussion

We have analyzed the range of values that *F*_*ST*_ can take as a function of the frequency *M* of the most frequent allele at a multiallelic locus, for an arbitrary value of the number of subpopulations *K*. We showed that *F*_*ST*_ can span the full unit interval only for a finite set of values of *M*, at 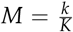 for integers *k* in [1, *K* − 1]. For all other *M, F*_*ST*_ necessarily lies below 1. The number of subpopulations *K* enlarges the range of values that *F*_*ST*_ can take as it increases.

This study provides the most complete relationship between *F*_*ST*_ and *M* obtained to date, generalizing previous results for the case of *K* = 2 subpopulations [12] and for a restriction to *I* = 2 alleles [14]. Interestingly, the maximal *F*_*ST*_ we have obtained merges patterns observed in these previous studies. Fixing *K* = 2, we obtain the upper bound on *F*_*ST*_ in terms of *M* that was reported by Jakobsson *et al*. [12]. As *K* increases, the piecewise pattern seen by Jakobsson *et al*. [12] for the maximal *F*_*ST*_ in the *K* = 2 case for *M* in 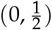 is observed in the multiallelic case for *M* in 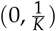. The decay from 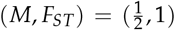 to (*M, F*_*ST*_) = (1, 0) seen by Jakobsson *et al*. [12] for *K* = 2 is observed for *M* in the decay from 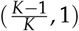 to (1, 0) for arbitrary *K*.

The allele frequency values for which the upper bound is reached for *M* in 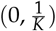 generalize those seen for the case of *K* = 2 and *M* in 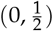 [12]. The upper bound is reached when all alleles are private, each subpopulation has as many alleles as possible at frequency *KM*, and at most one additional allele. The allele frequency values for which the upper bound is reached for *M* in 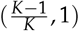 also generalize those seen for *K* = 2 and *M* in 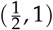: the maximum is reached when the most frequent allele is fixed in all subpopulations except one, and a single private allele is present in this last subpopulation.

The results from Alcala and Rosenberg [14] for *I* = 2 produce a more constrained upper bound on *F*_*ST*_ than for arbitrary *I*, with the domain of *M* restricted to 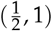. Nevertheless, many properties of the maximal *F*_*ST*_ we observe for unspecified *I* and *M* in 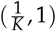 are similar to those seen for *I* = 2 and *M* in 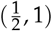: finitely many peaks at points 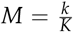, local minima between the peaks, and an increase in coverage of the unit square for (*M, F*_*ST*_) as *K* increases. The maximal *F*_*ST*_ functions for *M* in 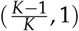 for unspecified *I* and for *I* = 2 agree, as the number of alleles required to maximize *F*_*ST*_ in this interval in the case of unspecified *I* is simply equal to 2.

In assuming that the number of alleles is unspecified, we found that the number of distinct alleles needed for achieving the maximal *F*_*ST*_ is 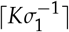 for *M* in 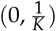 and *K* − ⌊*σ*_1_⌋ + 1 for non-integer *M* in 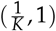; the maximum can be achieved with each number of distinct alleles in 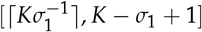, for *M* equal to 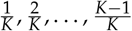. With a fixed maximal number of distinct alleles, such as in the *I* = 2 case of Alcala and Rosenberg [14] with *K* specified and in the *K* = 2 case with *I* specified [13], the upper bound on *F*_*ST*_ is less than or equal to that seen in the corresponding unspecified-*I* case. For *K* = 2, specifying *I* has a relatively small effect in reducing the maximal value of *F*_*ST*_ [13]. As in Edge and Rosenberg [13], specifying *I* in the case of larger values of *K* is expected to have the greatest impact on the *F*_*ST*_ upper bound at the lowest end of the domain for *M*.

In coalescent simulations, we found that the joint distribution of *M* and *F*_*ST*_ within their permissible space can help separate the impact of mutation and migration. Although the dependence of *F*_*ST*_ on mutation and migration rates has been long documented, the symmetric effects of mutation and migration under the island model [22] illustrate the difficulty in separating their effects. Under the island model, allele frequency *M* is informative about the scaled mutation rate 4*Nμ*, and comparing the value of *F*_*ST*_ to its maximum given *M* is informative about the scaled migration rate 4*Nm*. Adding a dimension that is more sensitive to mutation than to migration—*M* in our case— enables the separation of their effects. Other statistics, such as total heterozygosity *H*_*T*_ or within-subpopulation heterozygosity *H*_*S*_, have the potential to play a similar role [20].

Our results can inform data analyses. In particular, we caution users to examine upper bounds on *F*_*ST*_ to assess how mathematical constraints influence observations. As the constraints are strongest for *K* = 2, this step is valuable in pairwise comparisons; it is also useful when the frequency *M* of the most frequent allele can be small in relation to the number of populations *K*, such as for high-diversity forensic [28] and immunological [29] loci in human populations. Visual inspection of the values of *M* and *F*_*ST*_ within their bounds can suggest that constraints have an effect. 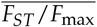 can provide a helpful summary by evaluating the proximity of *F*_*ST*_ values to their maxima.

Further, joint use of *M* along with *F*_*ST*_ could be useful in various applications of *F*_*ST*_, such as in inference of model parameters by approximate Bayesian computation [30] and machine-learning [31]. *F*_*ST*_ outlier tests to detect local adaptation from multiallelic loci [32] could search for *F*_*ST*_ values that represent outliers not in the distribution of *F*_*ST*_ values, but rather, outliers in relation to associated upper bounds. Computing null distributions for *F*_*ST*_ conditional on *M* could enhance the approach.

In an example data analysis, we have shown that taking into account mathematical constraints on *F*_*ST*_ can help understand puzzling *F*_*ST*_ behavior. In our example, *F*_*ST*_ at a set of loci was higher when comparing *K* = 6 chimpanzee populations than when comparing humans and chimpanzees (*K* = 2), even though the same loci were used and the mean value for *M* was similar in the two comparisons. A comparison of *F*_*ST*_ values to their respective maxima explained these counterintuitive results.

We note that analyses of *F*_*ST*_ in relation to *M* differ from analyses of *F*_*ST*_ in relation to within-subpopulation statistics *H*_*S*_ and *J*_*S*_ = 1 − *H*_*S*_, such as those performed in deriving the influential Hedrick’s 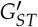 [9] and Jost’s *D* [33] statistics. We have previously shown that for biallelic loci in *K* subpopulations, for fixed *M*, the statistics 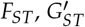, and *D* are all maximized at the same set of allele frequency values [15]. Although the normalizations of *F*_*ST*_ used to produce 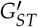 and *D* lead to statistics that are unconstrained in the unit interval as functions of 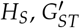 and *D* continue to be constrained as functions of *M*. A statistic that instead normalizes *F*_*ST*_ by its maximum as a function of *M*, a statistic of the total population, captures aspects of the allele-frequency dependence of *F*_*ST*_ that differ from those captured by normalizations by functions of within-subpopulation statistics.

In human populations, efforts to understand *F*_*ST*_ patterns trace in large part to Lewontin’s foundational *F*_*ST*_ -like variance-partitioning computation [34], in which it was seen that among-population differences (analogous to *F*_*ST*_) were small relative to within-population differences (analogous to 1 − *F*_*ST*_). Studies using loci with different numbers of alleles, loci with different frequencies for the most frequent allele, and samples with different numbers of subpopulations have varied to some extent in their numerical estimates of *F*_*ST*_ [14, 35, 36, 37, 38]. Mathematical results on *F*_*ST*_ bounds provide part of the explanation for these differences: they establish that each data set differing in the character of its loci and subpopulation set has its own distinctive interval in which its associated *F*_*ST*_ calculation could potentially land. Hence, each data set can give rise to a numerically distinct value not due to features of the underlying human biology, but rather, due to different constraints on the *F*_*ST*_ measure itself. *F*_*ST*_ bounds contribute to explaining quantitative variation in variance-partitioning computations—in which, although numerical values differ, the within-population component of genetic variation consistently predominates. The mathematics serves to support the qualitative claim that worldwide human genetic differentiation measurements represented by *F*_*ST*_ -like statistics have low values—as was argued by Lewontin fifty years ago.

## Data accessibility

All data are publicly available (see cited references).

## Authors’ contributions

NA and NAR designed the study. NA analysed the data. NA and NAR wrote the manuscript.

## Competing interests

Where authors are identified as personnel of the International Agency for Research on Cancer/World Health Organization, the authors alone are responsible for the views expressed in this article and they do not necessarily represent the decisions, policy or views of the International Agency for Research on Cancer/World Health Organization.

## Funding

Support was provided by NIH grant R01 HG005855, NSF grant BCS-2116322, and a France-Stanford Center for Inter-disciplinary Studies grant.

## Acknowledgements

We thank Kent Holsinger for comments on the manuscript and Maike Morrison for helpful conversations.

## Appendix Proof of eq. 3

This appendix derives the upper bound on *F*_*ST*_ as a function of *σ*_1_ (eq. 3). First, we separate the case of integer values of *σ*_1_ (eq. 3). First, we separate the case of *σ*_1_ integer values of *σ*_1_. Next, for non-integer values of *σ*_1_, we reduce the problem of maximizing *F*_*ST*_ to the problem of maximizing the sum of squared allele frequencies across alleles and subpopulations, 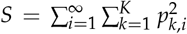. Next, we maximize *S* as a function of *σ*_1_, separately for *σ*_1_ in (0, 1) and for non-integer *σ*_1_ in (1, K).

### A useful expression for *F*_*ST*_

Suppose *K*≥ 2 is a specified integer. Suppose *σ*_1_ is a fixed value, with 0 < *σ*_1_ < *K*. We leave the number of alleles *I* unspecified. For each *i* ≥1, we write 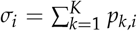, with *σ*_*i*_ ≥ *σ*_*j*_ for each *i* and *j* with *I* ≤ *j*. For convenience, *σ*_1_ is taken to mean both the function that computes the sum 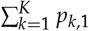 for a specified set of values of the *p*_*k,i*_ and a fixed value for that sum.

For each (*k, i*) with 1 ≤ *k* ≤ *K* and *i* ≥ 1, *p*_*k,i*_ lies in [0, 1], and 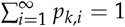 for all *k*, 1 ≤ *k* ≤ *K*. Define *F*_*ST*_ as in eq. 2. We seek to maximize *F*_*ST*_ over all possible sets of values of the *p*_*k,i*_ with a fixed value *σ*_1_ for the sum 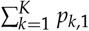. Note that because *σ*_1_ < *K* and 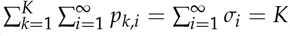, it follows that *σ*_2_ > 0.

Denote the sum of squared frequencies of allele 1 across subpopulations, 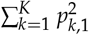, by *S*_1_. Denote 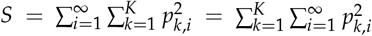 for the corresponding sum of squared frequencies of all alleles. We express eq. 2 in terms of *σ*_1_, *S*_1_, and *S*:

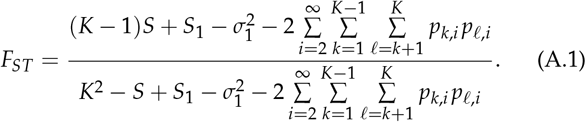

By construction of eq. 2, the denominator of eq. A.1 lies in (0, *K*^2^), as 0 < *H*_*T*_ < 1 from the fact that *σ*_2_ > 0. The numerator lies in [0, *K*^2^), as 0 ≤ *H*_*S*_ ≤ *H*_*T*_ < 1, so that 0 ≤ *H*_*T*_ − *H*_*S*_ < 1. *F*_*ST*_ lies in [0, 1], as 0 ≤ *H*_*S*_ and 0 < *H*_*T*_ imply 0 ≤ (*H*_*T*_ − *H*_*S*_)/*H*_*T*_ ≤ 1.

### The case of integer values of *σ*_1_

In eq. A.1, the numerator is less than or equal to the denominator, with equality if and only if 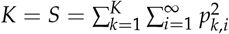. This equality in turn requires that for each *k*, there exists some *i* for which *p*_*k,i*_ = 1, a condition that can be achieved only if *σ*_1_ is an integer.

#### Theorem 1

*Suppose σ*_1_ *is an integer value*, 1, 2, …, *K*− 1. *F*_*ST*_ = 1 *if and only if (i) p*_*k*,1_ = 1 *in each of σ*_1_ *subpopulations, and (ii) for each of the K* − *σ*_1_ *remaining subpopulations, there exists a value of i* ≥ 2 *with p*_*k,i*_ = 1.

*Proof. F*_*ST*_ = 1 if and only if *S* = *K*, and *S* = *K* if and only if for each *k*, there exists an associated *i* with *p*_*k,i*_ = 1. For a fixed integer value of *σ*_1_, *p*_*k*,1_ = 1 in exactly *σ*_1_ subpopulations.□

Note that any set of equivalence relationships can exist among the values of *i* associated with the *K* −*σ*_1_ subpopulations in which *p*_*k*,1_ = 0, provided that none of these values of *i* is associated with more than *σ*_1_ subpopulations. For example, these values of *i* can be mutually distinct, or groups of them with size as large as *σ*_1_ can be mutually equal.

### Non-integer values of *σ*_1_

For non-integer *σ*_1_, the numerator of eq. A.1 is strictly less than the denominator. Hence, if the other quantities in eq. A.1 are fixed, then *F*_*ST*_ decreases with increasing 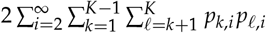 We have the following theorem.

#### Theorem 2

*Suppose σ*_1_ *is not an integer. F*_*ST*_ *satisfies*

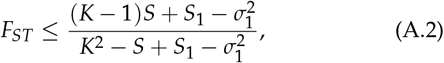

*equality requiring that for each i* ≥ 2, *there exists at most one value of k for which p*_*k,i*_ > 0.

*Proof*. Because 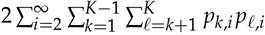 is subtracted in both the numerator and the denominator of eq. A.1, and because the numerator is strictly less than the denominator for non-integer *σ*_1_, *F*_*ST*_ can be bounded above by minimizing this term. Because *p*_*k,i*_ ≥ 0 for all (*k, i*), each sum 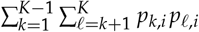 is bounded below by zero. Setting the sum to 0 for all *i* ≥ 2 gives the upper bound in eq. A.2.

For the equality condition, 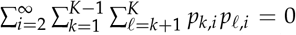 if and only if all products *p*_*k,i*_ *p*_*l,i*_ are zero—that is, if and only if for each *i* ≥ 2, at most one value of *k* has *p*_*k,i*_ > 0.□

By Theorem 2, to maximize *F*_*ST*_ for fixed non-integer *σ*_1_, we must maximize the quantity in eq. A.2. It suffices to consider sets of values of *p*_*k,i*_ in which for each *i* ≥2, at most one value of *k* has *p*_*k,i*_ > 0.

### The case of (non-integer) *σ*_1_ in (0, 1)

In this section, we find the set of values of the *p*_*k,i*_ that maximize *F*_*ST*_ for *σ*_1_ in (0, 1). We proceed in two steps. (i) We show that for *σ*_1_ in (0, 1), the maximal *F*_*ST*_ occurs at a set of *p* values for which *all* alleles are private: that is, for each *i* ≥ 1, *p*_*k,i*_ > 0 for at most one value of *k*. (ii) We determine the set of *p*_*k,i*_ values that, with all alleles private, maximizes *F*_*ST*_.

i. In eq. A.2, note that 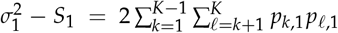. Because 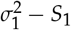 is subtracted from both numerator and denominator in eq. A.2, the quantity in eq. A.2 is maximal when 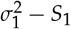 is minimal. In other words, the upper bound on *F*_*ST*_ is maximal if and only if 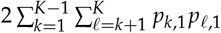 is minimal. Because *σ*_1_ < 1, a minimum of 0 for 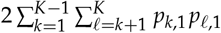 is achieved if and only if there is a single value *k* = *k′* at which *p*_*k*_*′* _,1_ = *σ*_1_, so that *p*_*k*,1_ = 0 for all *k* ≠ *k′*. We then have 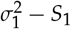, and from eq. A.2,

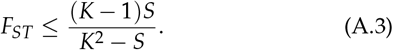 Each allele is private, and because allele 1 is the most frequent, *p*_*k,i*_ lies in [0, *σ*_1_] for all (*k, i*).
ii. The problem of finding the set of *p*_*k,i*_ values that maximizes *F*_*ST*_ has now been reduced to the problem of maximizing the right-hand side of eq. A.3, with the constraint that all alleles are private. Because the numerator in eq. A.3 increases with *S* and the denominator decreases with *S*, the maximum is achieved if and only if *S* achieves its maximal value. In other words, we seek to maximize 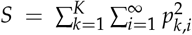, with the constraints 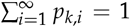 and *p*_*k,i*_ ≤ *σ*_1_ for each (*k, i*) with 1 ≤ *k* ≤ *K* and *i* ≥ 1. Because each allele is private, the maximum is achieved by separately maximizing each 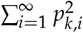 with constraints 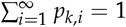 = 1 and *p*_*k,i*_ ≤ *σ*_1_.

This maximization is precisely that of Lemma 3 of Rosenberg and Jakobsson [39]. Applying the lemma, the maximum is achieved with *p*_*k*,1_ = *p*_*k*,2_ = … = *p*_*k,J*−1_ = *σ*_1_, *p*_*k,J*_ = 1 − (*J* − 1)*σ*_1_, and *p*_*k,i*_ = 0 for *i* > *J*, where 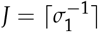 .It satisfies 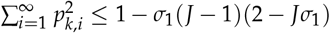. In other words, each subpopulation *k* possesses *J* − 1 private alleles with frequency *σ*_1_ and one private allele with frequency 1−(*J* − 1)*σ*_1_. Hence, *S* ≤ *K*[1 − *σ*_1_(*J* − 1)(2 − *Jσ*_1_)], so that eq. A.3 leads to eq. 3 for *σ*_1_ in (0, 1).

### The case of non-integer *σ*_1_ in (1, *K*)

This section finds the set of values of the *p*_*k,i*_ that maximizes *F*_*ST*_ for non-integer *σ*_1_ in (1, *K*). For non-integer 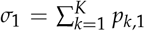 in (1, *K*), because 0 ≤ *p*_*k*,1_ ≤ 1 for all *k, p*_*k*,1_ > 0 for at least two values of *k*. Writing *S*\* = *S* − *S*_1_, Eq. A.2 can be rewritten

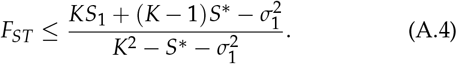

Because the numerator increases with *S*_1_, and because the numerator increases with *S*\* and the denominator decreases with *S*\*, the upper bound on *F*_*ST*_ is greatest when both *S*_1_ and *S*\* are maximized subject to 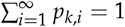 for each *k* and 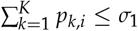 for each *i*. If *S*_*1*_ and *S*\* can be simultaneously maximized at the same set of values of the *p*_*k,i*_, then this set of values of the *p*_*k,i*_ achieves the maximal *F*_*ST*_.

We proceed in three steps. (i) First, we find the set of values of the *p*_*k,i*_ that maximizes *S*_1_. (ii) Next, we find the set of values that maximizes *S*\*. (iii) We then conclude that because the same set maximizes both *S*_1_ and *S*\* separately, this set achieves the upper bound in eq. A.4, and hence in eq. A.2.

i. We first maximize *S*_1_ for fixed non-integer *σ*_1_ in (1, *K*). More precisely, we seek to maximize 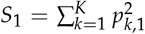 with constraints 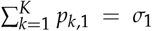 and *p*_*k*,1_ ≤ 1 for each *k* from 1 to *K*. This maximization is precisely that performed in Theorem 1 from Alcala and Rosenberg [14], a corollary of Lemma 3 of Rosenberg and Jakobsson [39]. Applying the theorem, the maximum is achieved by setting 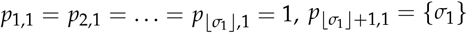, and *p*_*k*,1_ = 0 for all *k* > ⌊*σ*_1_⌋ + 1. The maximal value of *S*_1_ is {*σ*}^2^ + ⌊*σ*_1_⌋.
ii. Next, we maximize 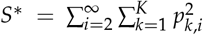 Because, by Theorem 2, all alleles with *i* ≥ 2 are private at the set of values of the *p*_*k,i*_ that maximizes *F*_ST_ for fixed non-integer *σ*_*1*_, each nonzero *p*_*k,i*_ for *i* ≥ 2 is equal to the associated *σ*_*i*_. The sum of the frequencies of all alleles across all subpopulations is 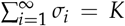, so that 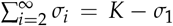. The problem of maximizing *S*^*\**^ is the problem of maximizing 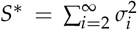 with the constraints 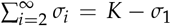 and *σ*_*i*_ ≤ 1 for each *i* from 2 to ∞. This maximization is again that performed in Lemma 3 of Rosenberg and Jakobsson [39]. Applying the lemma, the maximum is achieved by setting 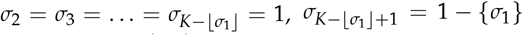, and *σ*_*i*_ = 0 for *i* > *K* –{⌊*σ*_1_⌋} + 1. The maximum is (1 − {⌊*σ*_1_⌋})^2^+ (*K* − ⌊*σ*_1_⌋ − 1).
iii. *S*_1_ is maximized at a set of *p*_*k,i*_ for which ⌊*σ*_1_⌋ subpopulations are fixed for allele 1, allele 1 has frequency{⌊*σ*_1_⌋}in one subpopulation, and allele 1 has frequency 0 in all other subpopulations. *S*\* is maximized at a set of *p*_*k,i*_ for which *K* − ⌊*σ*_1_⌋ − 1 subpopulations are fixed, each for a distinct allele *i* with *i* ≥ 2, one subpopulation possesses a distinct allele *i* ≥ 2 with frequency 1 − {*σ*_1_}, and all ⌊*σ*_1_⌋ other subpopulations possess no alleles *i* ≥ 2 of nonzero frequency.

The upper bound in eq. A.4 depends on both *S*_1_ and *S*\*, each of which depends on the *p*_*k,i*_. Were the set of values of the *p*_*k,i*_ that maximizes *S*_1_ and the set of values of the *p*_*k,i*_ that maximizes *S*\* to differ, additional work would be required to find the set of values of the *p*_*k,i*_ that maximizes *F*_*ST*_. However, we now observe that *S*_1_ and *S*\* can be simultaneously maximized at the same set of values of *p*_*k,i*_, so that the same set of values of the *p*_*k,i*_ maximizes *S*_1_ and *S*\* and hence *F*_*ST*_. In particular, ⌊*σ*_1_⌋ subpopu-lations are fixed for allele 1, each of *K* − ⌊*σ*_1_⌋ − 1 subpopulations is fixed for its own private allele, and a single subpopulation possesses allele 1 with frequency{*σ*_1_}and a private allele with frequency 1 − {*σ*_1_}. The number of alleles of nonzero frequency is *K−* ⌊*σ*_1_⌋ + 1. Only the most frequent allele is shared by more than one subpopulation, and a single subpopulation possesses more than one allele of nonzero frequency.

Substituting the maximal values of *S*_1_ and *S*\* into eq. A.4, for non-integer *σ*_1_ in (1, *K*), we obtain the maximal *F*_*ST*_ in terms of *σ*_1_ shown in eq. 3.

---

**Figure S1.**
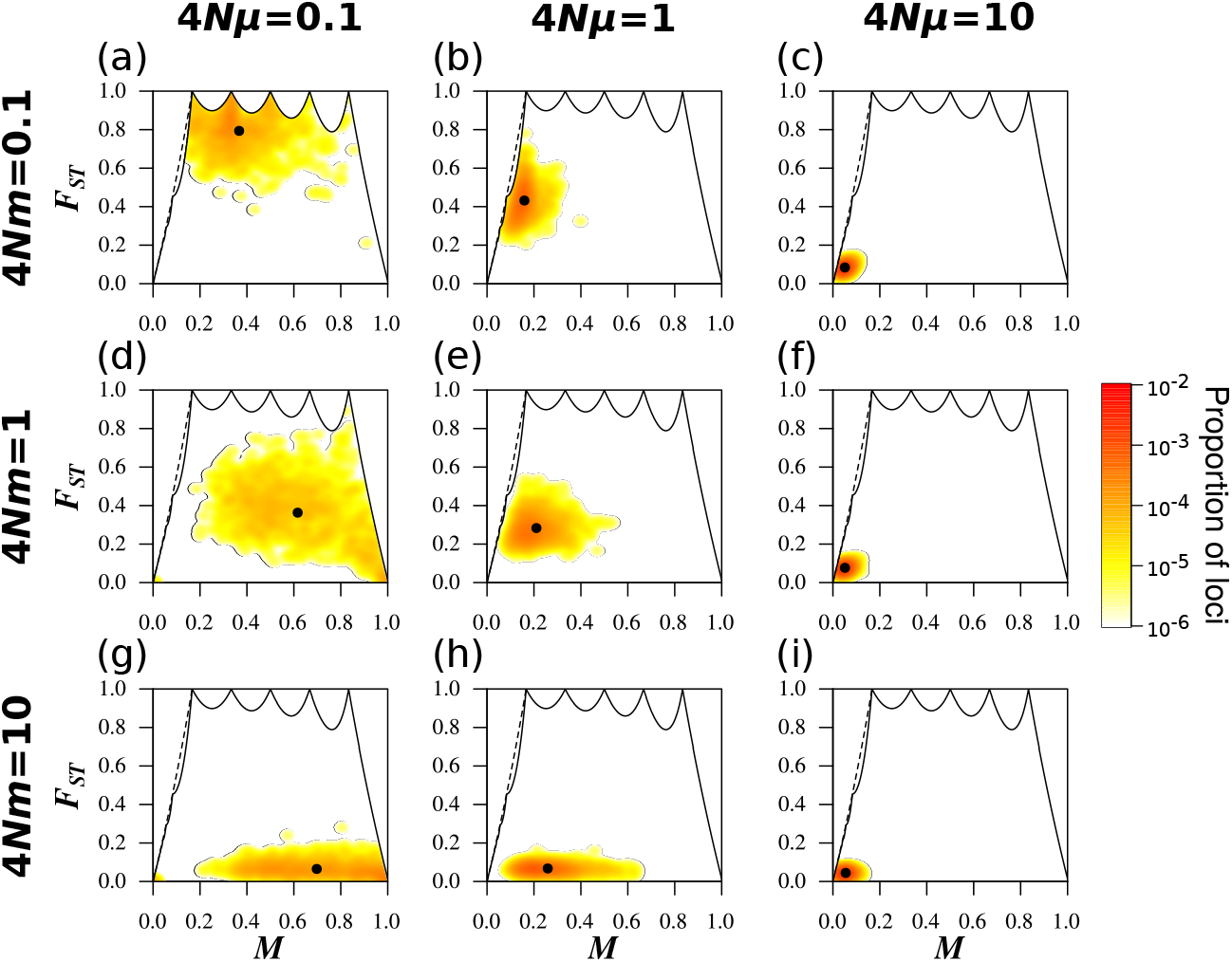
Joint density of the frequency *M* of the most frequent allele and *F*_*ST*_ in the island migration model with *K* = 6 subpopulations, for different scaled migration rates 4*Nm* and mutation rates 4*Nμ*. The simulation procedure and figure design follow Figure 2.

**Figure S2.**
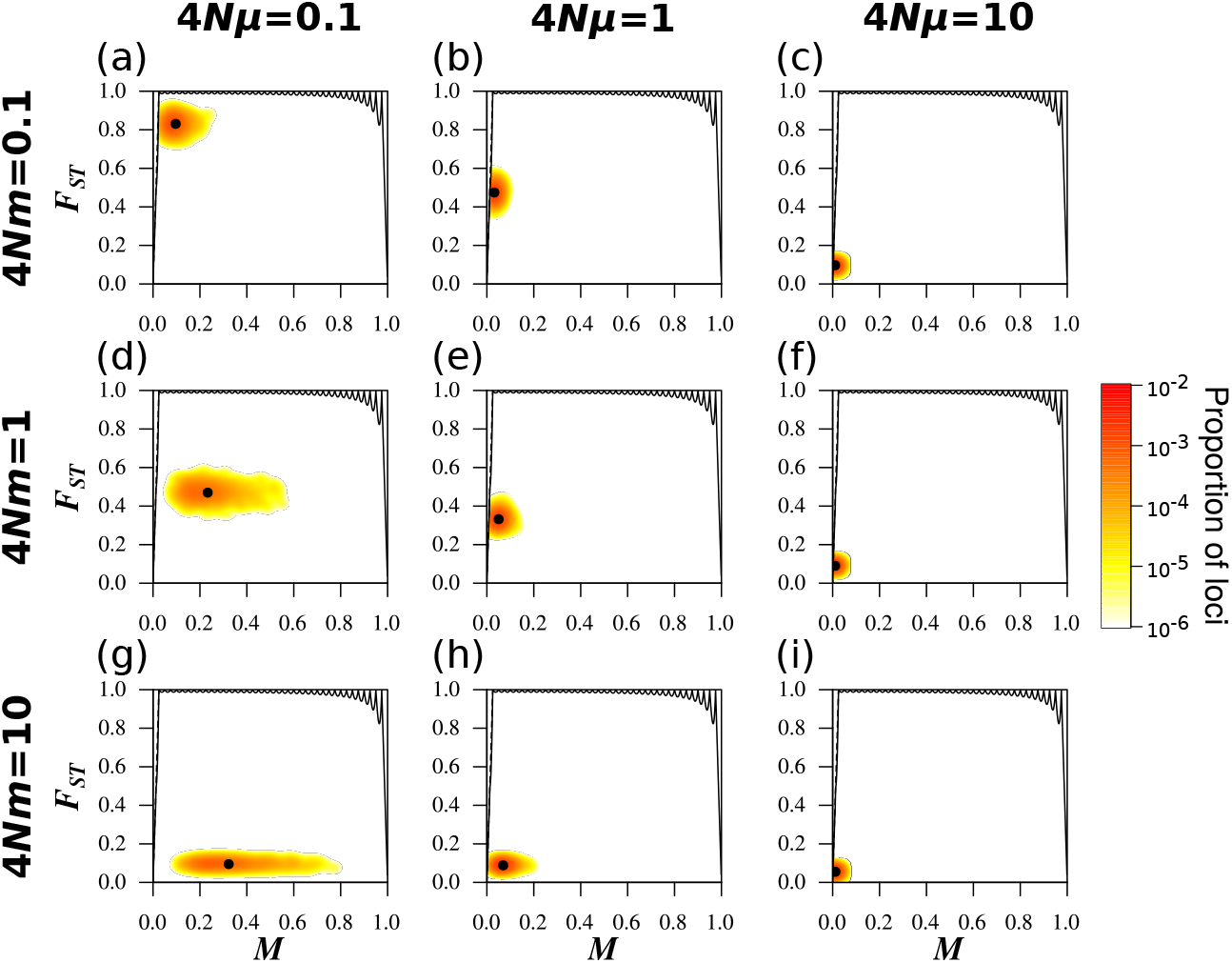
Joint density of the frequency *M* of the most frequent allele and *F*_*ST*_ in the island migration model with *K* = 40 subpopulations, for different scaled migration rates 4*Nm* and mutation rates 4*Nμ*. The simulation procedure and figure design follow Figure 2.

## Supplementary File S1: MS commands

We applied `MS`, specifying the scaled mutation and migration parameters. We performed the simulations for *K* = 2, *K* = 6, and *K* = 40 subpopulations. For each command, we replace x by the desired 4*Nμ* value and y by the desired 4*Nm* value.

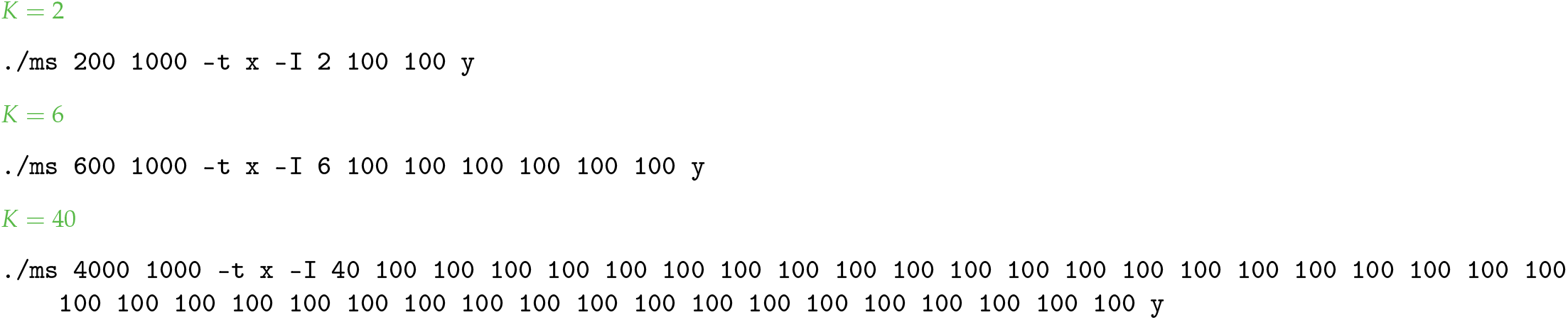

## Notes

### Competing Interest Statement

The authors have declared no competing interest.

### Summary of Updates

Correction of an omission in the conditions under which the upper bound on FST is reached, in the case of integer values of \sigma_1. This led to changes in the text and appendix primarily on pages 3 and 9, but also on pages 4 and 10. None of the figures changed. Additionally, additional citations and more explicit explanation of the purpose of the data application to Human and Chimpanzee data.

